# Enrichment of neural crest cells by antibody labelling and flow cytometry for single-cell transcriptomics in a lizard

**DOI:** 10.1101/2025.05.21.655068

**Authors:** Robin Pranter, Cedric Patthey, Nathalie Feiner

## Abstract

Neural crest cells (NCCs) are a key component of the vertebrate body plan and contribute to a variety of different traits. However, their dynamic migratory behavior and spatiotemporal heterogeneity in the developing embryo pose significant challenges for their identification and isolation. Consequently, most studies of NCCs have been confined to model organisms with established transgenic tools. To overcome this limitation, we present a novel approach that combines antibody labelling with fluorescence activated cell sorting to enrich for NCCs and we demonstrate the approach in the common wall lizard (*Podarcis muralis*). Through microscopy, reverse transcription quantitative polymerase chain reaction and single-cell RNA sequencing, we show that the method enriches for NCCs as efficiently as methods relying on transgenic animals. Using this technique, we successfully characterise transcriptional profiles of NCCs in wall lizard embryos. We anticipate that this method can be applied to a wide range of vertebrates that lack transgenic tools, enabling deeper insights into the roles that neural crest cells are playing in developmental and evolution.

## Introduction

Neural crest cells (NCCs) are multipotent stem cells of the embryo with the unique ability to generate derivatives typically associated with all three germ layers. As such, NCCs give rise to a multitude of different cell types such as chromatophores in the epidermis, neurons and glia of the peripheral nervous system, chondrocytes and osteocytes that make up the facial skeleton, and several types of endocrinal cells such as the chromaffin cells of the adrenal gland, important for behavioural aspects. While much has been learned about the diversification of neural crest cells into a variety of cell types within a handful of model organisms, it remains poorly understood how the regulation of NCCs has been diversified throughout the evolution of vertebrates to give rise to diverse pigmentation, cranial morphologies and behaviours. Beyond understanding the diversification of developmental processes, from an evolutionary perspective it would be interesting to assess if the developmental linkage between certain traits imposed by their common origin in NCCs may lead to co-evolution of these traits. For example, NCCs have been suggested to be important for the evolution of several phenotypic syndromes that have been observed in wild and domesticated animals (Brandon et al., 2023; Feiner et al., 2024; Wilkins et al., 2014).

To understand how NCCs themselves have evolved, and how they may have affected the evolution and potentially co-evolution of NCC-derived traits, it will be necessary to study the development of NCCs in a broad range of taxa. The current gold standard for studying cellular diversity and differentiation is single cell transcriptomics (scRNA-seq). However, there are several challenges associated with scRNA-seq of NCCs and their derivatives. First, NCCs are migratory and dispersed among other cells, which means that they cannot be isolated through (micro-)dissection. Second, the ability of NCCs to differentiate into a diverse set of cell types, many resembling cells of other embryonic origins, makes it challenging to label NCCs using genetic markers. Early markers will not be expressed in all derived lineages and late markers will be shared with phenotypically similar cells descending from other stem cells. To overcome these challenges, scRNA-seq studies of NCCs have relied on established transgenic lines of mice (*Mus musculus*) and zebrafish (*Danio rerio*) that express permanently a reporter following expression of a marker gene for (a subset of) NCCs. For example, cells expressing *Sox10, Wnt1* or *Pax2*, have been targeted using fluorescence activated cell sorting (FACS) (Howard et al., 2021; Soldatov et al., 2019; Xu et al., 2024). Alternatively, studies have used chicken embryos (*Gallus gallus*), which are accessible for manipulation during development, and have employed injection or electroporation of various cell-labelling constructs such as plasmids or viruses (Jacobs-Li et al., 2023; Morrison et al., 2017; Williams et al., 2019). However, many of the taxa that would be candidates for investigating evo-devo questions about NCCs do not have established methods for transgenesis or other manipulations during development. Furthermore, the reproductive biology of many species may pose practical challenges, for example by restricting access to embryos to a particular time of year, necessitating methods for storing of embryos or cells. To date, many workflows in scRNA-seq are optimized for fresh cells, and where these are not available, the only viable alternative is to resort to single nuclei RNA-seq since isolation of nuclei is feasible from fixed or stored tissue (Habib et al., 2017; Habib et al., 2016).

To overcome these challenges and make the isolation of NCCs available to non-model organisms, we present a methodology based on antibody labelling of fixed cells followed by FACS and demonstrate its use in the common wall lizard (*Podarcis muralis*). This species is interesting in this context because NCCs have been implicated in the evolution of a suite of correlated traits (i.e., a phenotypic syndrome), including coloration, cranial morphology and behaviour (Feiner et al., 2024). Our previous work has characterized the dynamic distribution of NCCs in this species and found that immunohistochemistry (IHC) targeting the epitope Human Natural Killer 1 (HNK-1) is a reliable marker of NCCs (Pranter & Feiner, 2024), similar to what has been found for chicken (Tucker et al., 1988) and other squamate reptiles (veiled chameleon: Diaz Jr et al., 2019; Egyptian cobra: Khannoon et al., 2021; California kingsnake: Reyes et al., 2010; brown anole: Weberling et al., 2025). Common wall lizards can be bred in captivity, but the breeding season is restricted to a few months per year, during which females lay 2- 3 clutches with about 2-10 eggs per clutch (Böhme, 1986). Thus, access to embryos is restricted to a short period, which makes the ability to store cells advantageous.

Our methodology consists of the following steps (see Fig. 1): 1) fixation and long-term storage of dissociated cells in methanol, 2) staining NCCs with HNK-1, and 3) enriching for the HNK-1-labelled NCCs using flow cytometry. The approach is similar to previously established methods that target intracellular molecules, for example transcription factors, instead of surface molecules, to enrich for certain cell types (Hrvatin et al., 2014; Pan et al., 2011), which is commonly used in immunological studies (Albu et al., 2010; Cossarizza et al., 2017). We validate the efficiency of our protocol using a combination of microscopy, reverse transcription quantitative polymerase chain reaction (RT-qPCR) and scRNA-seq, and use the latter to gain insight into the transcriptional profile of NCCs in the common wall lizard.

**Figure 1.**
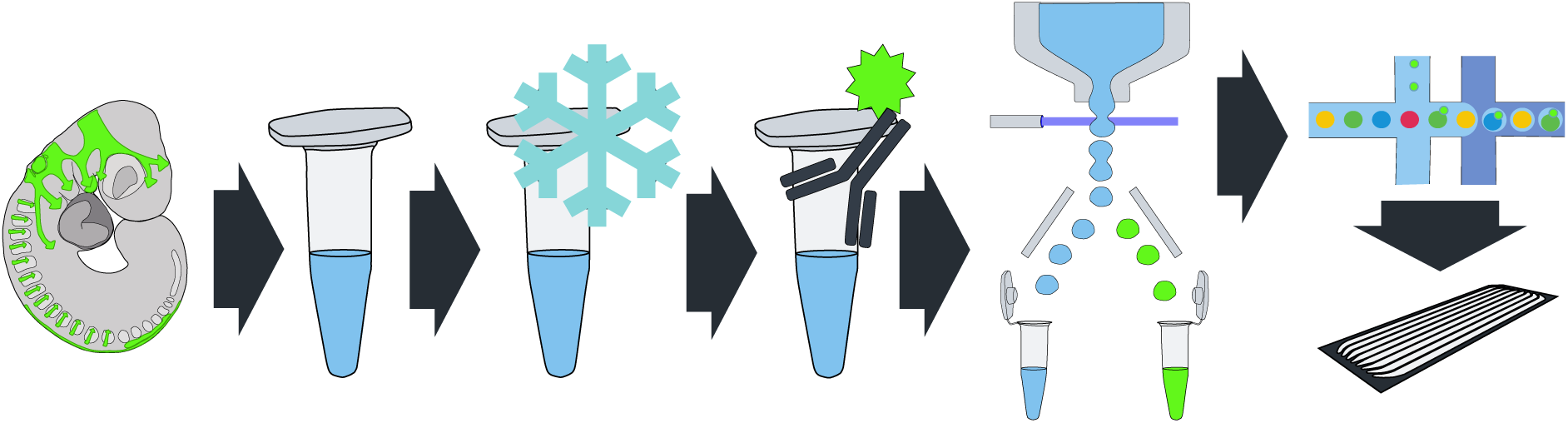
Workflow for enrichment of NCCs by HNK-1 labelling and flow cytometry. NCCs (shown in bright green; see Pranter & Feiner, 2024) and other embryonic cells (shown in grey) are dissociated into single cells, fixed and stored in methanol at -80°C. A fluorophore-conjugated HNK-1 antibody is used to target NCCs during flow cytometry. HNK-1-positive, NCC-enriched cells are subjected to library preparation and scRNA-seq. See Material and Methods and detailed protocol in the supplementary material.

## Results

In brief, our experiments resulted in a method starting with eggs collected from a captive colony of common wall lizards that were dissected to obtain embryos. The embryos were dissociated into cell suspensions using trypsin and fixed in methanol. The dissociated and fixed cells were stored at -80°C. To enrich for NCCs, cell suspensions were stained using the antiHNK-1 antibody and FAC-sorted based on the HNK-1 fluorescence signal. RNA was preserved during antibody staining using a MOPS-based buffer supplemented with RNAse inhibitors and DTT as previously described (Patthey et al., 2016). For a detailed protocol, see supplementary material. In the following sections we describe each step and its validation.

### Staining NCCs with an antibody binding the epitope Human Natural Killer 1 (HNK-1)

NCC-staining by the HNK-1 antibody has previously been confirmed in *P. muralis* (Pranter & Feiner, 2024), albeit with paraformaldehyde- and not methanol-fixed embryos. To rule out the possibility that methanol fixation affects the HNK-1 epitope and thereby prevents efficient labelling of NCCs fixed in methanol, we performed immunohistochemistry staining of HNK-1 using methanol-fixed embryos. The embryo was also stained with 4′,6-diamidino-2-phenylindole (DAPI) to provide spatial context in the embryo. This resulted in successful labelling of NCCs, evidenced by HNK-1 signal in migrating NCCs entering the second and third pharyngeal arches in the head and passing through the somites in the trunk. Some differentiation can be seen in the peripheral nervous system such as the trigeminal nerve, Fig. 2), which is expected at this somite stage (Pranter & Feiner, 2024).

**Figure 2.**
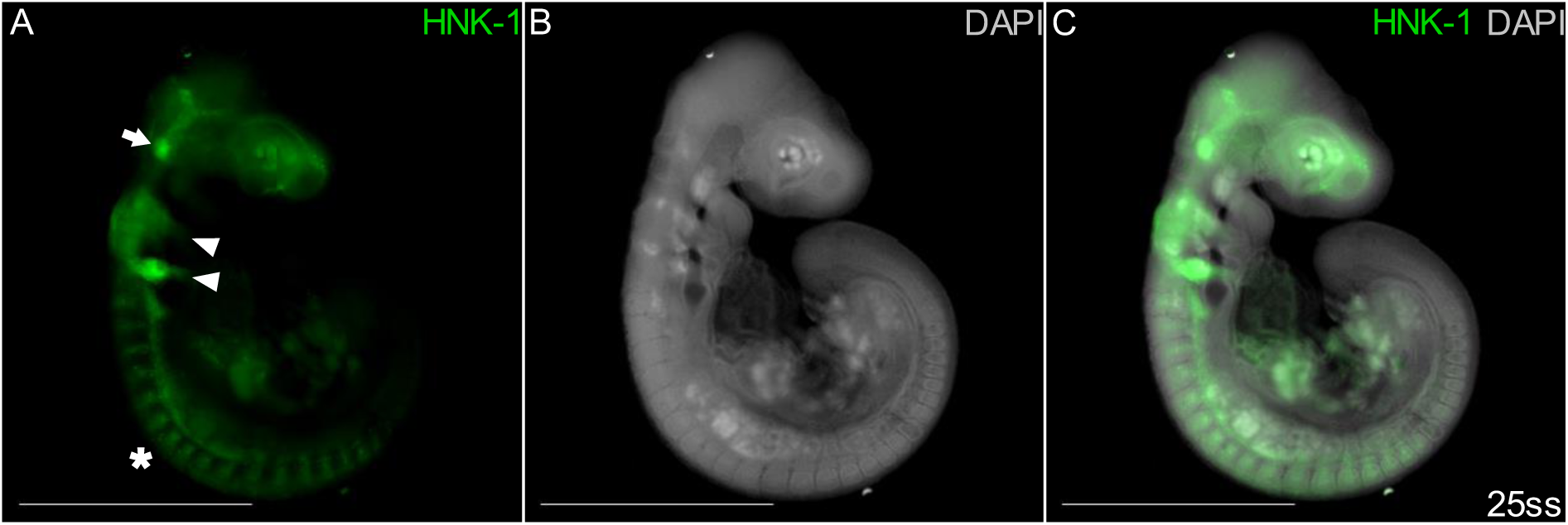
Migratory and differentiating NCCs are stained by HNK-1 in a methanol-fixed embryo at somite stage 25. **A** HNK-1 distinctly labels cranial NCCs in migratory streams entering the second and third pharyngeal arches (arrowheads), multiple streams of migratory trunk NCCs (asterisk) and differentiating peripheral nervous system (e.g. the trigeminal nerve; arrow). **B-C** DAPI staining visualizes regions with high cellular density. Scale bars are 1 mm.

### Enriching for the HNK-1-labelled NCCs using flow cytometry

To obtain suspensions of dissociated, fixed cells, whole embryos were minced and further dissociated using trypsin and filtered through a cell strainer. Cells were subjected to a fixable Live/Dead stain to label dead cells and to efficiently identify high-quality cells using flow cytometry (see below). The cells in the resulting suspension were fixed using methanol and stored for up to several month at -80°C (also in methanol) prior to staining with DAPI and fluorophore (FITC) conjugated anti-HNK-1, and FACS.

To sort HNK1-positive single NCC cells that were alive at the onset of fixation, the cells were selected using a series of nested gates (i.e. sub-setting criteria) (Fig. 3). Debris and doublets were excluded based on forward- and side-scatter properties (Fig. 3A-C) as well as DAPI intensity, selecting for cells with DAPI intensity characteristic of nuclei in G1 and G2 phase of the cell cycle (Supplementary figure 1). Among the selected events (corresponding to cells), variation in size and shape (forward- and side-scatter) was continuous (Fig. 3A-C). The integrity of cell morphology was also confirmed using ocular examination of sorted cells using a light microscope, and no severe signs of cell leakiness were detected (Supplementary figure 2). The anti-HNK-1 (FITC) signal was distributed as one large cluster of FITC-negative cells followed by a long tail of cells with gradually more FITC-signal (Fig. 3E), and samples that were not stained with HNK-1 (FITC) lacked this tail with high FITC signal (Fig. 3F). This profile was consistent across experiments (Fig. 3G) and shows similarity to flow cytometry experiments sorting NCCs using GFP signal in transgenic embryos (Howard et al., 2021; Jacobs-Li et al., 2023).

**Figure 3.**
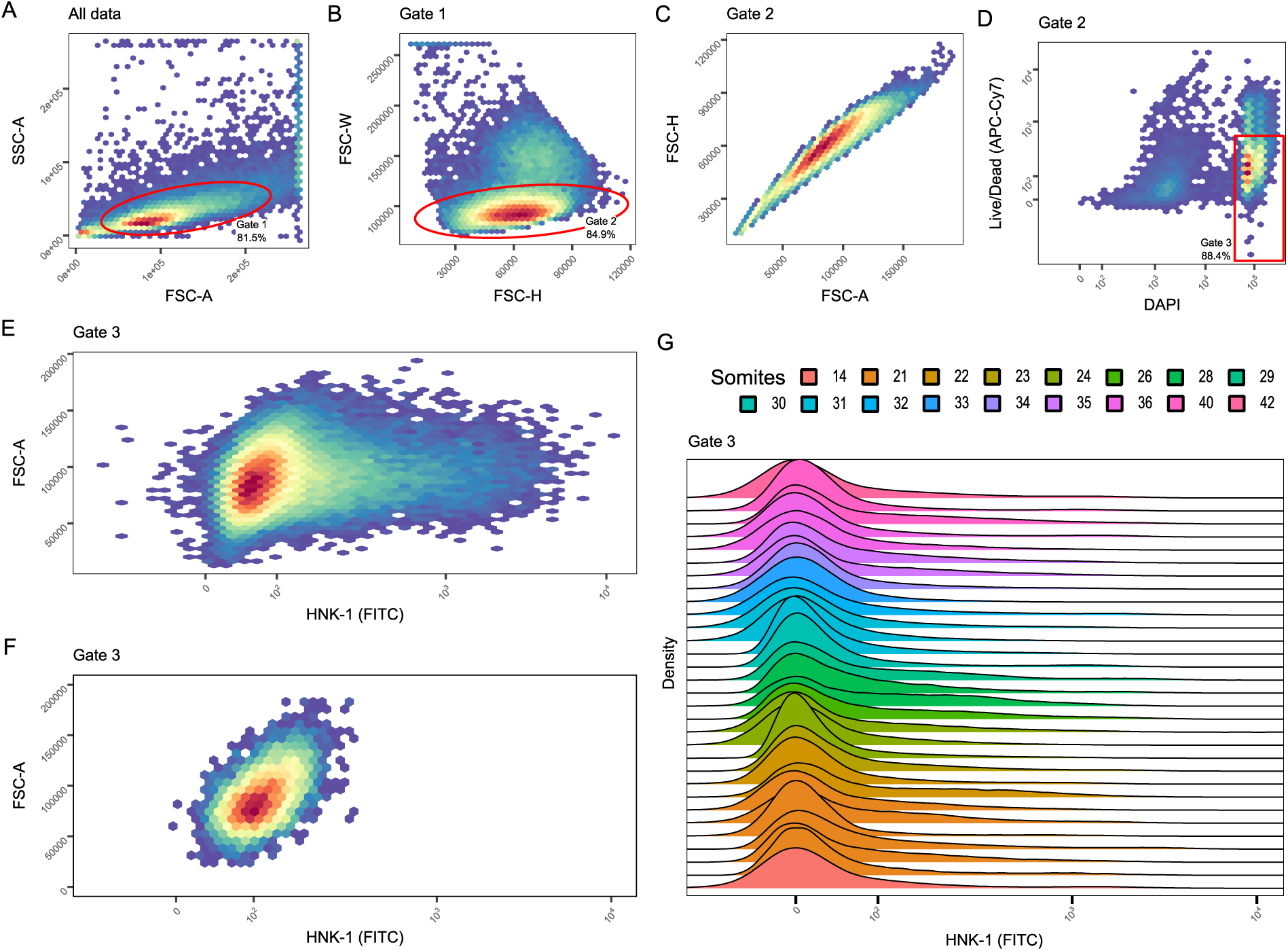
Fluorescence-activated cell sorting of HNK-1 positive cells. Panels A-F show heatmaps of cell densities of one representative sample, while panel G summarizes flow cytometry results of 31 samples. **A** Forward and side scatter are plotted for all cells. Gate 1 selects for proper cells and against debris. **B** Height and width of the forward scatter plotted for the cells in gate 1. Gate 2 selects for single cells and against doublets. **C** Area and height of the forward scatter confirms that no doublets are contained in gate 2. **D** Fluorescence from the DAPI and Live/Dead stains plotted for the cells in gate 2. Gate 3 selects the DAPI positive and Live/Dead negative cells, (i.e., cells with DNA that were alive before fixation). **E** FITC fluorescence plotted against forward scatter area of cells in gate 3. A long tail of cells with gradually more FITC fluorescence can be seen emerging from the main cluster with low FITC fluorescence. **F** Same plot as shown in panel E, but for a sample that was not stained with HNK-1 (negative control) that shows the lack of the tail of FITC-positive cells. **G** The distributions of FITC signals in 31 samples. The peak of each distribution has been centered to align the distrubutions with highest FITC signal at 0 for the purpose of visualization. Overall profiles with a large peak or FITC-negative cells followed by a long tail of gradually more FITC-positive cells, are similar across experiments.

### Characterization of HNK-1 positive cells using RT-qPCR

To assess whether HNK-1 positive cells obtained through FACS were enriched for NCCs, we collected different sets of cells from flow cytometry and subjected them to RT-qPCR targeting the three NCC marker genes *Sox10, FoxD3* and *Snai2*, whose utility as NCC-markers have been previously confirmed in the common wall lizard (Pranter & Feiner, 2024). First, RNA integrity in fixed, frozen and sorted cells was confirmed by analysing RNA extracts using Bioanalyzer (RIN: mean = 7.7, sd = 0.99, n = 4) and found to be comparable to cells that were sorted directly after fixation (RIN = 6.4, n = 1). Two pools, each consisting of three embryos to ensure sufficiently high cell numbers, were stained and gated as described above and further sorted into bins with increasingly stronger HNK-1 (FITC) signal using four additional gates (visualized in Fig. 4A): the first gate included cells below the 90^th^ percentile of the distribution (‘Gate Neg’), the second one cells between the 90^th^ and the 95^th^ percentiles (‘Gate 90%’), the third one cells between the 95^th^ and the 97.5^th^ percentiles (‘Gate 95%’) and the fourth one cells above the 97.5^th^ percentile (’Gate 97.5%’). RT-qPCR of the resulting four bins showed that the relative expression level of each of the three marker genes *Sox10, FoxD3* and *Snai2* was positively correlated with the HNK-1 (FITC) signal (Fig. 4B-D, table 1). *Sox10* and *FoxD3* showed the highest expression levels in the 97.5^th^ percentile bin with a marked, gradual increase from leniently to strictly sorted samples with an increasing HNK-1 (FITC) signal. The results for *Snai2* were less pronounced, but a trend towards higher expression levels with increased HNK-1 (FITC) signal was also evident. In sum, the RT-qPCR results of *Sox10* and *FoxD3* strongly support on enrichment of NCCs in the fraction of cells with high HNK-1 (FITC) signal.

**Figure 4.**
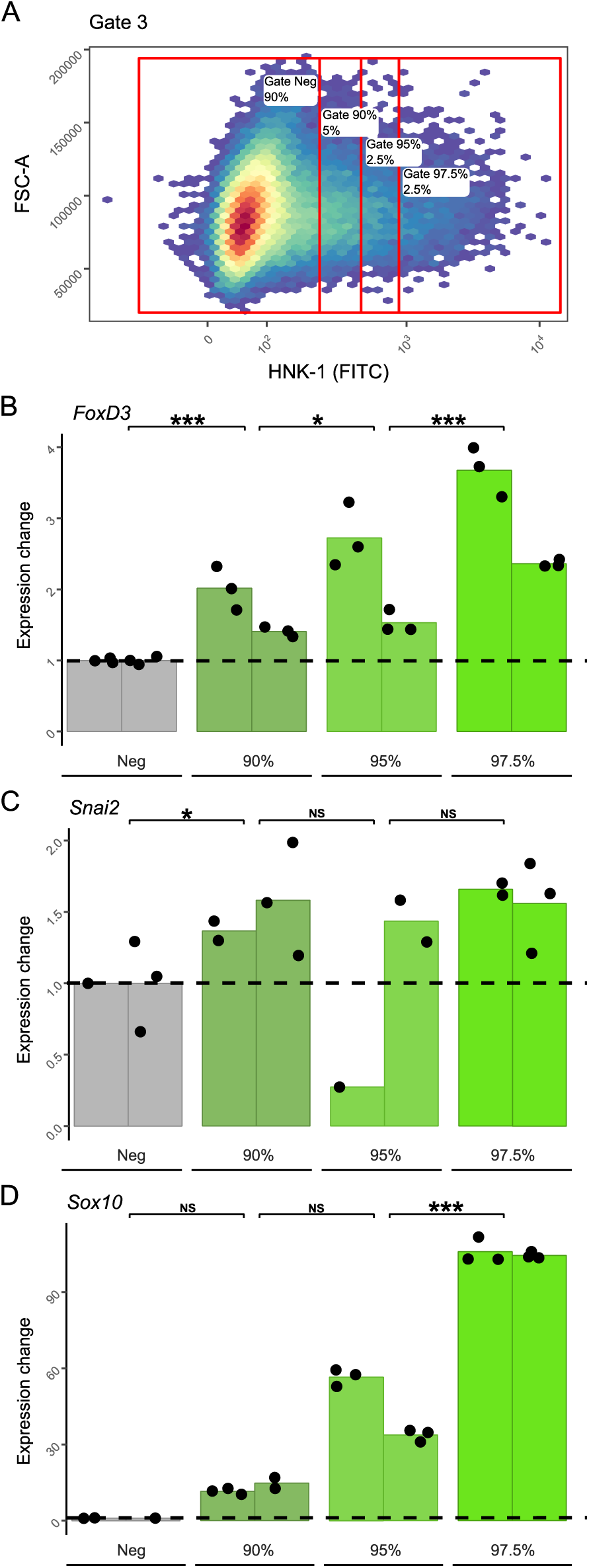
RT-qPCR confirms gradual enrichment of NCC-marker gene expression with increasing strength of HNK-1 signal in flow cytometry. **A** Cells from one of the two pools, gate 3 plotted as HNK-1 (FITC) signal against the forward-scatter area. The cells were further sorted using four new gates based on their FITC fluorescence (i.e., HNK-1 signal). The first gate includes cells below the 90^th^ percentile of the distribution (‘Gate Neg’), the second one cells between the 90^th^ and the 95^th^ percentiles (‘Gate 90%’), the third one cells between the 95^th^ and the 97.5^th^ percentiles (‘Gate 95%’) and the fourth one cells above the 97.5^th^ percentile (’Gate 97.5%’). (**B-D**) Normalized expression divided by the ‘Gate Neg’ mean normalized expression of the expression levels of the NCC markers *FoxD3, Snai2* and *Sox10* measured by RT-qPCR. Technical replicates are plotted as points and the mean of each biological replicate (pool) is plotted with one bar per sorting gate. Statistical significance of a positive relationship between NCC marker expression and FITC signal is indicated by asterisks.

**Table 1.** NCC marker expression increases with increasing HNK-1 (FITC) signal. Resulting contrasts for linear models of relative gene expression as a function of gate averaged across biological replicates. The analysis is run as one model per marker and *P*-values are Tukey adjusted.

| *Marker* | *Contrast* | *Estimate* | *SE* | *df* | *t ratio* | *P-value* |
| --- | --- | --- | --- | --- | --- | --- |
| *FoxD3* | Neg - 90% | -0.00451 | 0.000915 | 19 | -4.933 | 0.0001 |
|  | 90% - 95% | -0.00224 | 0.000915 | 19 | -2.453 | 0.024 |
|  | 95% - 97.5% | -0.00677 | 0.000915 | 19 | -7.403 | <.0001 |
| *Snai2* | Neg - 90% | -0.000163 | 6.93E-05 | 12 | -2.344 | 0.0371 |
|  | 90% - 95% | 0.000119 | 7.51E-05 | 12 | 1.58 | 0.1402 |
|  | 95% - 97.5% | -0.000142 | 7.51E-05 | 12 | -1.894 | 0.0826 |
| *Sox10* | Neg - 90% | -0.00607 | 0.0209 | 15 | -0.29 | 0.7757 |
|  | 90% - 95% | -0.01575 | 0.0174 | 15 | -0.907 | 0.3789 |
|  | 95% - 97.5% | -0.05826 | 0.0165 | 15 | -3.526 | 0.0031 |

### Characterization of HNK-1 positive cells using scRNA-seq

To further validate the enrichment of NCCs using HNK-1 labelling combined with FACS, we generated scRNA-seq datasets from unsorted and unstained cells, cells sorted above the 85^th^ percentile of HNK-1 (FITC) fluorescence, or cells sorted above the 95^th^ percentile of HNK-1 fluorescence. The unsorted and the leniently sorted samples were pools of each three embryos and were subjected to the 10X Chromium protocol. The strictly sorted sample was derived from a single embryo and was subjected to the Smart- seq3 protocol (replicated in two 384-well plates) since this method is more suitable for samples with low numbers of cells, which result from the stricter sorting cut-off, as starting material (Hagemann-Jensen et al., 2020).

Comparison between the unsorted and the leniently sorted samples (both sequenced using the same methodology) confirms that staining and FAC-sorting does not influence standard quality metrics for scRNA-seq data such as the gene and read counts per cell and the percentage of mitochondrial counts per cell (Fig. 5, Supplementary table 1). The strictly sorted sample shows higher gene and read counts per cell, which is due to different scRNA-seq protocols, but comparable metrics in the percentage of mitochondrial counts per cell. The percentage ribosomal counts appears to decrease with stricter sorting.

**Figure 5.**
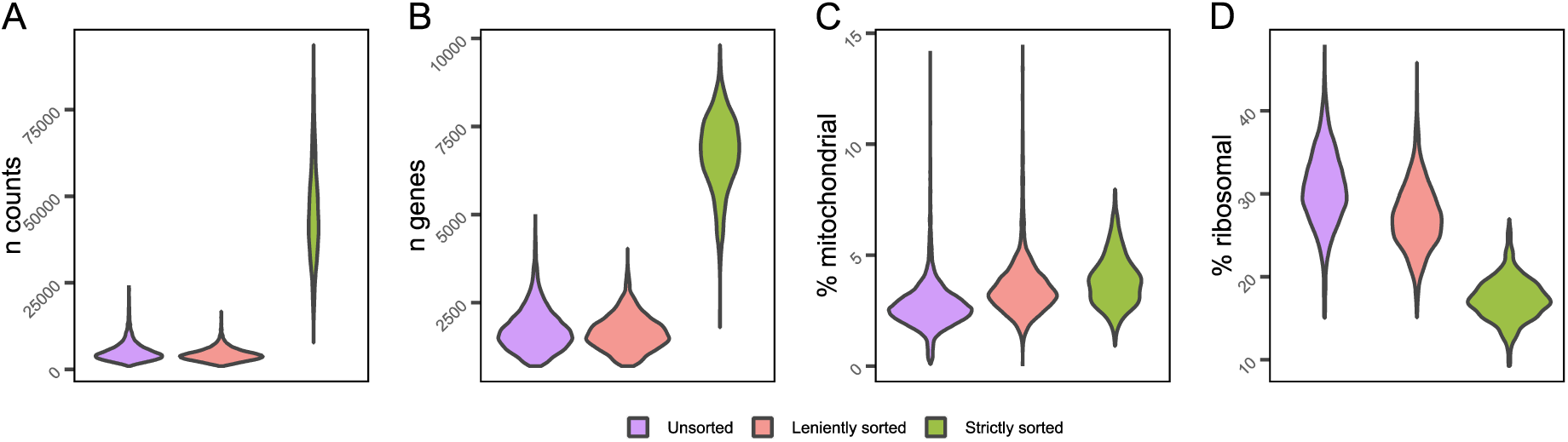
Quality metrics of the scRNA-seq datasets. Quality metrics for scRNA-seq data plotted for each sample. The two strictly sorted and the unsorted samples are each pooled together, resulting in 10674 unsorted, 3036 leniently sorted and 694 strictly sorted cells. The values plotted on the y-axes are: **A** per cell total counts of unique molecular identifiers (UMI). **B** per cell total gene count, **C** the per cell percentage of UMIs that are derived from mitochondrial genes and **D** the per cell percentage of UMIs that come from ribosomal genes.

We employed three tests to assess the efficiency of our approach to enrich for NCCs: 1) comparison of mean NCC-signature scores calculated from marker gene lists between unsorted and sorted samples; 2) projection of each sample onto a reference mouse whole-embryo atlas and qualitative assessment of cell types; 3) calculation of relative expression of the NCC marker gene *Sox10* and comparing the percentage of *Sox10*-expressing cells between unsorted and sorted samples of this and published datasets.

First, NCC enrichment was assessed by calculating the NCC-signature score of each cell and comparing median scores between the three different types of samples. The NCC- signature score was calculated using a core set of NCC marker genes (Simoes-Costa & Bronner, 2015): *Sox10*, *FoxD3*, *Snai2*, *Pax3, Pax7, Sox9, Wnt1, Tfap2a, Sox5* and *Zic1*. The NCC-signature score overall provided strong support for an enrichment of NCCs in the strictly sorted sample. The median NCC-signature score for the strictly sorted sample was 0.144, while the equivalent number in leniently sorted sample was -0.026 and for the unsorted sample, it was -0.026. There were significant differences between all samples (Wilcoxon rank sum tests: unsorted vs leniently sorted, *P*-value = 0.006; unsorted vs strictly sorted, *P*-value < 0.001; leniently sorted vs strictly sorted, *P*-value < 0.001).

Second, to classify cell types, cells from each sample were projected onto a reference dataset consisting of the mouse whole-embryo atlas (Cao et al., 2019) using the scmap method (Kiselev et al., 2018), resulting in a cell type classification for each cell in the dataset. The cells were reclassified as ‘NCCs’ (including derivatives), ‘possible NCCs’ or ‘not NCCs’ according to established knowledge of which cell types are derived from NCCs (Barresi & Gilbert, 2021; Bronner & LeDouarin, 2012; Cao et al., 2019; Eames et al., 2020; Le Douarin & Kalcheim, 1999). The ‘possible NCCs’ include cell types that are derived by both NCCs and other stem cells, for example chondrocytes. In the unsorted sample, 55.0% of cells were classified as either NCCs or possible NCCs, which was lower than in the reference (62.1%; Cao et al., 2019). In the leniently sorted sample this percentage was 64.7% (Supplementary figure 3) and in the strictly sorted sample the corresponding value is 74.7% (the projection of the latter is shown in Fig. 6A). Performing the same projection using a published dataset derived from transgenic mice that was enriched for NCCs (Soldatov et al., 2019), returned 96.1% of cells classified as NCCs or possible NCCs (Fig. 6B).

**Figure 6.**
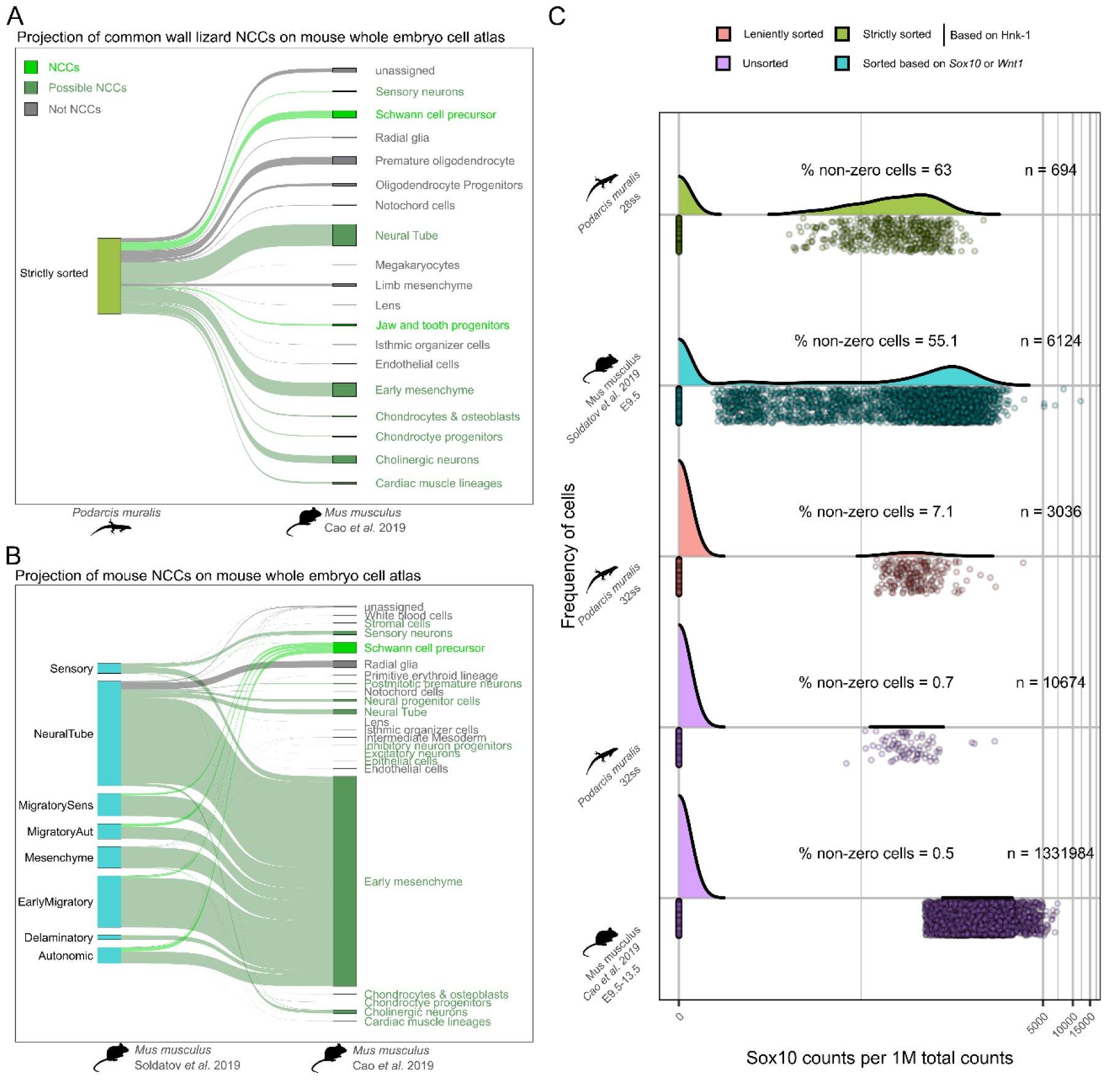
Comparison of HNK1-labelled and FAC-sorted cells of the common wall lizards to published datasets of mouse. **A** Projection of the strictly sorted samples onto the whole embryo mouse cell atlas. **B** Projection of a mouse NCC dataset (Soldatov et al., 2019) onto the whole embryo mouse cell atlas (Cao et al., 2019). The NCCs are labelled on the left by their published cell type labels. **C** Relative expression of *Sox10* of each sample (the strictly sorted and unsorted samples are each pooled in this plot) as well as mouse sorted NCCs (Soldatov et al., 2019) and whole embryo cells (Cao et al., 2019). The total number of cells and the percentage of non-zero cells are given above each curve.

Third, we assessed the efficiency of our method by calculating the relative expression of *Sox10* per cell as the number of counts per million reads and compared the distribution of expression values between our samples and two previously published datasets from mouse derived from whole embryos (Cao et al., 2019) or transgenesis-mediated, sorted NCCs (Soldatov et al., 2019). Across all datasets, the distribution of relative *Sox10* expression per cell showed a bimodal pattern with cells either expressing no *Sox10* at all or expressing it at moderate to high levels (Fig. 6C). We therefore considered it justified to compare the proportion of cells with non-zero *Sox10* expression across datasets. Using this metric, we found that the whole embryo datasets were comparable between lizard (0.7%) and mouse (0.5%). This metric of the proportion of *Sox10* expressing cells was slightly increased in the leniently sorted sample with 7.1% and was considerably higher in the strictly sorted sample of the common wall lizard with 62.3%. The corresponding metric in the published mouse NCC dataset (Soldatov et al., 2019) was slightly lower with 55.1%. Considering the insights gained from the three independent tests, we conclude that our methodology using HNK-1 labelling coupled with FAC-sorting enriches for NCCs as efficiently as sorting based on transgenic animals.

### Differential expression analysis in unsorted cells reveals novel NCC markers

To gain further insight into NCC gene expression profiles, the unsorted samples that allow us to contrast NCCs and non-NCCs were examined closer. All cells with a NCC signature score higher than 0.15 were defined as NCCs. In a Uniform Manifold Approximation and Projection (UMAP) projection, NCCs were found to distribute among non-NCCs in much, but not all, of transcription-space (Fig. 7A). Assessing differential gene expression between NCCs and all other cells identified 597 genes with an NCC-related expression (Fig 7B-C, Supplementary data table 1). 27% of the NCC markers were found to be previously mentioned together with the phrase ‘neural crest’ in a PubMed search.

**Figure 7.**
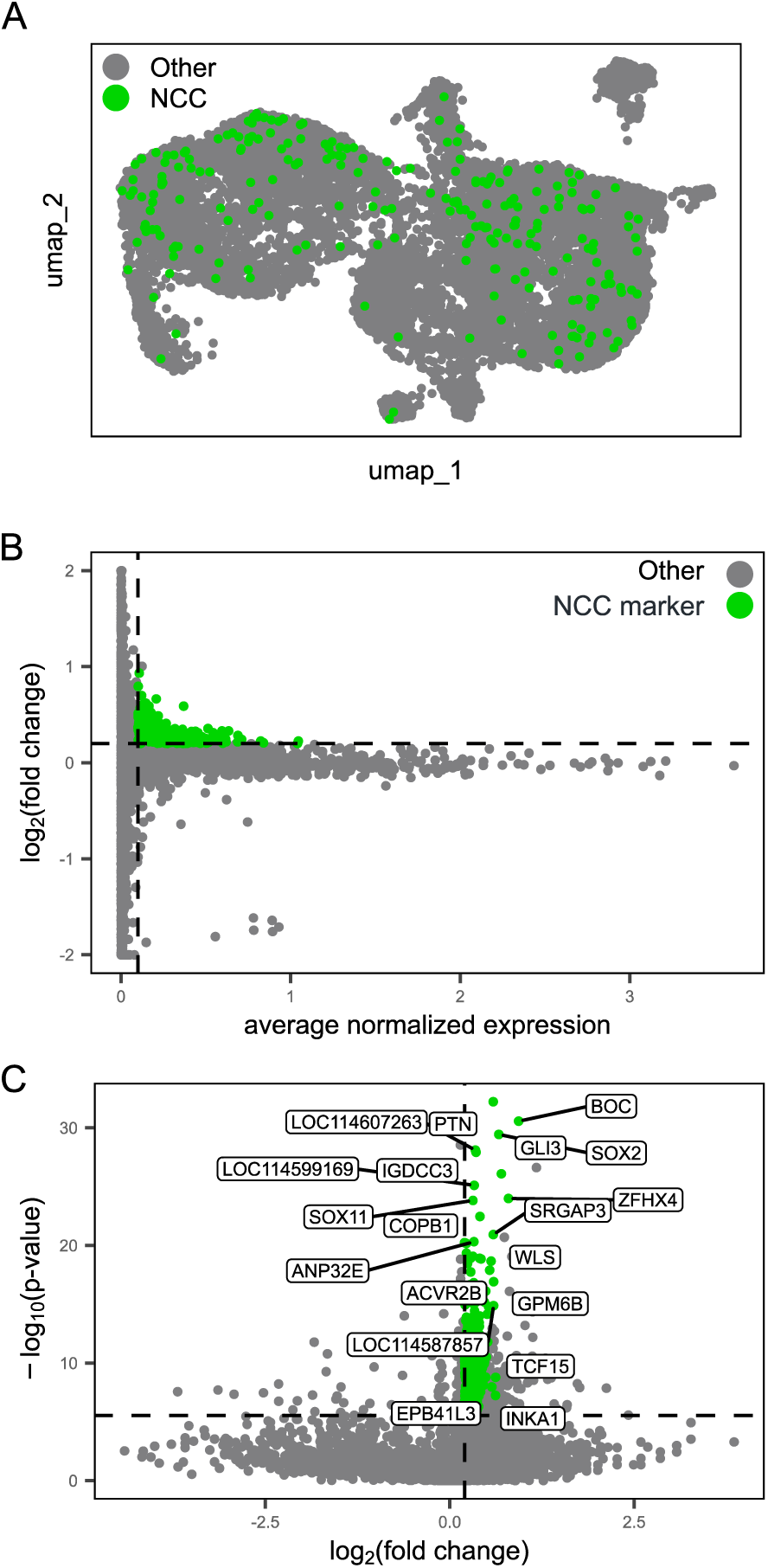
Transcriptional profile of NCCs and unbiased identification of novel NCC marker genes. **A** UMAP projection of all unsorted cells coloured according to their classification as putative NCCs (green) or others (grey; 232 out of 10674 cells had an NCC signature score above 0.15 and were classified as putative NCCs). Putative NCCs do not form a uniform cluster, but distribute among other cells in UMAP space. **B-C** 597 out of 17830 genes were defined as positively differentially expressed in putative NCCs compared to other cells. Thresholds marked by dashed lines (average normalized expression > 0.05, log_2_ fold change > 0.1 and Bonferroni corrected p value < 0.05). The genes with the largest and/or most significant differences that had meaningful names are labelled in **C**.

## Discussion

To understand how neural crest cells have diversified across vertebrates, and how their developmental organization has shaped the evolution of NCC-derived cells and traits, novel methods for targeting NCCs in non-traditional model organisms are needed. To meet this demand, we developed a method for NCC isolation that does not rely on established transgenic lines. We also aimed to develop a method that can be applied to organisms that pose challenges associated with obtaining embryos and rapidly processing fresh samples on demand. To achieve this, we used the common wall lizard, in which a syndrome composed of neural-crest derived traits has evolved (Feiner et al., 2024). We combined methanol fixation that allows long-term storage of embryonic cells at -80°C with labelling of NCCs with the antibody HNK-1 followed by flow cytometry. We confirmed that the resulting cells collected from embryonic common wall lizards were of high quality, contained RNA that was not degraded, and were enriched in NCCs to the same extent as published data from a transgenic mouse model.

Only a small fraction of all embryonic cells originates from the neural crest. Due to their dynamic nature and partial resemblance to cells of other embryonic origin, it is difficult to count the exact number of NCCs or neural crest-derived cells in an embryo. However, the expression of NCC marker genes such as *Sox10* in whole embryo single-cell profiling datasets that we report here suggests that NCCs make up less than 1% of all cells in an embryo. This illustrates that an enrichment strategy is necessary for a targeted study of this cell type. Our flow cytometry results showed that HNK-1 labelled cells do not form two separated populations but rather a continuous distribution, reminiscent of results obtained using transgenic animals (Howard et al., 2021; Jacobs-Li et al., 2023). The reason for this may be that NCCs exhibit varying levels of HNK-1 on their surface, and even non-NCCs, such as certain blood cells, may have HNK-1 molecules on their surface (Abo & Balch, 1981). Nevertheless, using a series of gradually stricter gating, we were able to gradually enrich our sample for NCCs. Strictly sorting samples by taking only the top ∼5% of cells with the highest HNK-1 (FITC) signal proved to satisfactorily enrich for NCCs and increased the proportion from below 1% to over 60% of isolated cells. This enrichment level is comparable to that attained by a published study using transgenic mice (Soldatov et al., 2019). When leniently sorting samples by taking the top ∼15% of cells, the enrichment level dropped to below 10%, suggesting that NCCs are strongly concentrated in the positive tail of the HNK-1 distribution.

One possible extension of the method we describe here is that different antibodies can be implemented. Using a single marker to target NCCs is not likely to capture all cells belonging to the NCC lineage. This is due to the challenging biology of NCCs with a high potency and the co-option of cellular phenotypes otherwise expressed by cells derived from other stem cells. Since HNK-1 mainly labels migratory NCCs, post-migratory NCCs will likely be missed particularly at later stages of development. It appears that there is no single marker that is both *specific* enough to *only* label NCCs and their derivatives, yet *sensitive* enough to label *all* NCCs including their derivatives. This is a challenge faced both by transgenesis-dependent and -independent methods. However, the flexibility of antibody guided FACS opens the possibility of combining different antibodies. Carefully selecting a panel of different antibodies should make it possible to enrich for a wider or more narrow range of cells and could be a further development of our methodology.

In addition to describing a new methodology, our study contributes the first list of unbiased NCC markers in a reptile. The list contains several genes with described functions in NCCs, for example *Mycn* (Hanemaaijer et al., 2021; Jansky et al., 2021; Simoes-Costa & Bronner, 2015), *Ctnnb1* (Javali et al., 2020; Simoes-Costa & Bronner, 2015) and *Pdgfra* (Cebra-Thomas et al., 2007; Soldatov et al., 2019), and also genes that have no known function in NCC biology. Studies of NCCs in reptiles typically rely on marker genes described in model organisms, yet this prevents the discovery of genes that may be expressed in NCCs in only some groups of organisms. While this list of genes should be considered preliminary and non-exhaustive due to the relatively shallow sequencing, it can be a valuable resource for further studies of NCC biology in reptiles and beyond.

In conclusion, we provide a new method for NCC-enrichment using anti-HNK-1 guided flow cytometry. The value of this method is that it enables the study of NCCs using scRNA-seq in all species where NCCs can be labelled by HNK-1 antibodies, and without the need for establishing transgenic tools. We anticipate that this will be important for future efforts into understanding the evo-devo of NCCs.

## Material and Methods

### Embryo collection

A captive colony of breading common wall lizards, wild caught in 2018 in central Italy, was kept at Lund University (see Feiner et al., 2018a for details on housing; 2018b). Breeding groups consisted of one male and one or two females per cage, and each cage had a pot of moist sand where females laid their eggs. The eggs were collected within 24 h of oviposition. One embryo from each clutch was immediately dissected and assigned a developmental stage based on somite counts, which was taken to represent the whole clutch (Feiner et al., 2018a, 2018b). The rest of the clutch was incubated at 24°C in moist vermiculate until they reached the desired developmental stage. Somite formation has previously been found to keep a constant rate of four somites per day when incubated at 24°C (Feiner et al., 2018a, 2018b).

### Immunohistochemistry of MeOH fixed embryos

Eggs were dissected in cold phosphate buffered saline (PBS; 10 mmol/L phosphate buffer, 2.7 mmol/L KCl, and 137 mmol/L NaCl, pH 7.4) and transferred to gradually higher concentrations of methanol in PBS containing 0.1% Tween20 and finally stored in methanol at -20°C. Whole-mount embryos were stained using immunohistochemistry with the primary antibody HNK-1 (1:500; Anti-Hu CD57 eBioscience 11-057742) following the method described in Pranter and Feiner (2024) and imaged using a Zeiss Axio Imager M2 florescence microscope. All reagents are listed in Supplementary table 2.

### Cell dissociation, fixation, and staining

Embryos were dissected from their eggs in nuclease-free PBS. Each embryo was photographed and then minced into fine pieces using a sterile scalpel. The minced embryo was incubated at 37°C with Trypsin (Gibco^TM^ TrypLE^TM^ Express Enzyme (1X), phenol red) to dissociate the cells into a single cell suspension. Trypsin treatment was terminated after ten minutes by applying an inhibitor (Gibco^TM^ Defined Trypsin Inhibitor). The suspension was triturated 50 times up and down through a 1ml pipet tip and then passed through a 70μm cell strainer (Flowmi^TM^ Cell Strainer) to remove any remaining clumps. To stain the cells for viability the suspension was incubated for 12 minutes on ice with a Live/Dead stain (Invitrogen LIVE/DEAD^TM^ Fixable Near-IR Dead Cell Stain Kit; L34975). Suspended cells were washed once in nuclease-free PBS, fixed in methanol and stored in methanol at -80°C for up to several months.

The fixed cell solutions were resuspended in a permeabilization buffer (PB; 2% BSA, 100U/ml RNAse-inhibitor, 5mM 1,4-Ditiotreitol, 0.1M MOPS pH 7.5, 1mM EGTA, 2mM MgSO4, 125mM NaCl and 0.1% Saponin), and incubated on ice for 5min. Once permeabilized, the cells were stained with a mix of DAPI (1:1000; ThermoScientific 62248), which stains DNA, and a FITC-conjugated HNK-1 antibody (1:100; Anti-Hu CD57 eBioscience 11-057742) in MRDB (a buffer identical to PB but lacking saponin), followed by a washing step with the same buffer and passed through a 70μm cell strainer.

### Flow cytometry and FACS

Dissociated cells were sorted in a FACS ARIA III machine. Debris, doublets and cells that were dead before fixation were removed by three gates (Fig. 3A-D). The first gates excluded debris by excluding cells with low values in both forward and side scatter area (Fig. 3A). The second gate removes doublets by excluding cells with high values in forward scatter width, which can be clearly made out when forward scatter width and height are plotted against each other (Fig. 3B). The third gate further removes additional (large) debris, non-nucleated cells and cells that were dead before fixation by only selecting cells with high DAPI fluorescence and low Live/Dead fluorescence (Fig. 3D). The remaining cells were inspected and sorted based on their anti-HNK-1 (FITC) signal (Fig. 3E).

### RT-qPCR

Cells dissociated, fixed and stained as described above and then FAC-sorted into 200µL of MRD buffer (identical to PB but lacking saponin and BSA) following the gating described above. An appropriate volume was moved to a new tube based on the known concentration of cells. Total RNA was extracted using an RNeasy Micro Kit including DNase digestion on the membrane and stored in nuclease free water at -80°C. RNA was reverse transcribed into cDNA using SuperScript III Reverse Transcriptase and oligo-dT primers according to the manufacturer’s instructions. The resulting cDNA was used as template for qPCR of *FoxD3*, *Snai2* and *Sox10*. The gene *GAPDH* which has an expected constant expression was analysed together with the three NCC-markers and was used as a reference in the normalization procedure. See Supplementary table 3 for primer sequences. Technical replicates were filtered based on their melting-curves to exclude incorrect amplification such as primer dimers, and the resulting Cq values were normalized as:

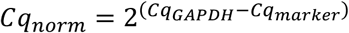

Differences in normalized Cq values between samples collected from different FACS gates were modelled linearly with normalized Cq as the dependent and biological replicate and FACS gate as factors. Least square means were estimated pairwise between each gate and its immediate neighbour(s) and p-values were Tukey corrected using the ‘emmeans’ package in R (Lenth, 2024).

### Single cell transcriptomic analysis

#### 10X Chromium

We used the 10X Chromium protocol for unsorted and leniently sorted samples. Cell-dissociations from 12 embryos with the average somite stage 32 were grouped into four pools. Two pools were put together from unsorted cells directly from -80°C storage. Two pools were put together from FAC-sorted samples following the staining and gating described above with a final sorting gate taking the cells above the ∼85^th^ percentile of the FITC distribution. The collected cells were pelleted by centrifugation, resuspended in methanol and stored at -80°C for four days before cell capture, and library preparation at the Eukaryotic Single Cell Genomics Facility, which is part of SciLifeLabs Sweden. Libraries were sequenced on Novaseq flow cells. The pools were processed into cDNA using 10X Chromium Next GEM Single Cell 3’ Reagent Kits v3.1 (Dual Index) and sequenced on an Illumina NovaSeq 6000. The target was ∼5000 cells per sample and ∼20k read pairs per cell. Reads were mapped to the PodMur_1.0 reference genome (Andrade et al., 2019) and gene by cell count matrices were generated using the Cellranger pipeline from 10XGenomics (Zheng et al., 2017) and further analysed using the R package Seurat 5.2.1 (Hao et al., 2024).

The count matrix was filtered in several steps. First, genes that were discovered in less than 3 cells, and cells that had less than 700 expressed genes, were removed. Second, cells whose transcriptome consisted of more than 15% mitochondrial reads were removed to exclude leaky cells and cells with less than 15% ribosomal reads were removed to exclude outliers. Doublets were predicted using DoubletFinder V2.0 (McGinnis et al., 2019) and removed from the dataset. Lastly, cells with more than 25000 reads were removed to exclude outliers. Expression levels were then normalized using SCTransform (Choudhary & Satija, 2022; Hafemeister & Satija, 2019) and variation associated with the percentage of mitochondrial reads and cell cycle state were removed by regression. The latter was modelled using the expression of orthologs of human markers for either the S-phase (38 genes) or the G- and M-phases (37 genes) (Tirosh et al., 2016). After filtering and normalization, the read count, gene count, percentage mitochondrial counts, percentage ribosomal counts and cell cycle scores did not appear to associate with the distribution of cells in reduced dimension space (PCA, tSNE and UMAP) or differ between predicted clusters.

#### Smart-seq3

Because cell counts in flow cytometry indicated that the number of cells obtained through strict sorting would be too few to allow application of the 10X Chromium method (in particular for early embryos), we used the Smart-seq3 protocol for strictly sorted samples. A cell dissociation from a 28ss embryo was stained and FAC-sorted as described above with a final sorting gate taking the cells above the ∼95^th^ percentile. The cells were sorted directly onto two 384 well plates using the ‘single-cell’ mode on the same FACS Aria III as described above. Plates were submitted for library preparation and sequencing to the Eukaryotic Single Cell Genomics Facility at SciLifeLab in Solna, Sweden. cDNA was prepared in each well following the Smart-seq3 protocol (Hagemann-Jensen et al., 2020) and sequenced on an Illumina NovaSeq XPlus. Reads were mapped to the PodMur_1.0 reference genome (Andrade et al., 2019) and the count matrices were generated using the zUMIs pipeline (Parekh et al., 2018). Count matrices were further analysed using the Seurat 5.2.1 package in R (Hao et al., 2024). The count matrix was filtered to exclude cells with less than 1000 expressed genes, more than 8% mitochondrial reads and/or less than 86% exonic reads. The two filtered count matrices were normalized using the SCTranform() function in Seurat (Choudhary & Satija, 2022; Hafemeister & Satija, 2019) and variation associated with percentage mitochondrial reads, and cell cycle state was regressed out. After filtering and normalization, the read count, gene count, percentage mitochondrial counts, percentage ribosomal counts and cell cycle scores did not appear to associate with the distribution of cells in reduced dimension space (PCA, tSNE and UMAP) or differ between predicted clusters.

#### Assessing NCC enrichment

We employed three different approaches to assess the efficiency of the enrichment for NCCs, and describe each of them here in turn.

To derive a NCC signature score per cell, we used the function AddModuleScore in the Seurat package with the following gene set: *Pax3, Pax7, FoxD3, Sox10, Sox9, Wnt1, Tfap2A, Sox5, Snai2* and *Zic1* (Simoes-Costa & Bronner, 2015). We then calculated the NCC signature scores for all cells in a given dataset and compared their mean across datasets.

To assess the proportion of cells expressing a core NCC marker gene, *Sox10*, we calculate the proportion of cells with non-zero expression of *Sox10* from each filtered count matrix and from the count matrix of published dataset from mouse (Cao et al., 2019; Soldatov et al., 2019). This was done using this formula:

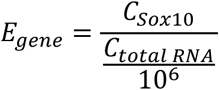

Where 𝐶_𝑔𝑒𝑛𝑒_ is the number of reads assigned to *Sox10* per cell, 𝐶_𝑡𝑜𝑡𝑎𝑙 𝑅𝑁𝐴_ is the total read count per cell, and 𝐸_𝑔𝑒𝑛𝑒_ is the relative expression of *Sox10* per cell. From this, the proportion of cells with non-zero *Sox10* expression was derived for each sample.

To derive a classification of cells into cell types, the count matrices were independently projected onto a reference mouse whole embryo cell atlas (Cao et al., 2019) using the scmapCluster() function in the scmap 1.28 package (Kiselev et al., 2018) using 1500 variable features and a likelihood threshold of 0.03. Gene orthology was assigned using an orthology table downloaded from Ensembl and only one-to-one orthologues were retained. For comparison, a dataset with sorted NCCs from mouse embryos (Soldatov et al., 2019) was also projected using the same likelihood threshold and number of variable features.

#### Constructing UMAP space and deriving an unbiased list of NCC marker genes

By selecting the cells from the unsorted samples that had NCC signature scores above 0.15, we divided the cells into putative NCCs and other cells. An unbiased list of NCC marker genes was derived from a differential gene expression analysis contrasting putative NCCs and other cells using the Seurat function FindMarkers. As criteria for calling genes differentially expressed, we applied the following: average normalized expression > 0.05, log2 fold change > 0.1 and Bonferroni corrected *P*-value < 0.05.

## Acknowledgements

We thank Anna Fossum at the Biomedical Center, Lund University, for assistance with fluorescence activated cell sorting, Nikolay Oskolkov at the National Bioinformatics Infrastructure Sweden at SciLifeLab for advice on bioinformatic analyses, Michael Bok for guidance in imaging, Tyra Westelius for help with embryo collection and Tobias Uller for comments on the manuscript. We acknowledge support from the National Genomics Infrastructure in Stockholm funded by Science for Life Laboratory, the Knut and Alice Wallenberg Foundation and the Swedish Research Council, and NAISS for assistance with massively parallel sequencing and access to the UPPMAX computational infrastructure.

## Data availability statement

All sequence data generated in this study have been deposited in NCBI with accession numbers XXX (*information will be provided upon acceptance*).

