## Supplementary Information for "Enrichment of neural crest cells by antibody labelling and flow cytometry for single-cell transcriptomics in a lizard"

This document contains:

-3 supplementary figures

-2 supplementary tables

-supplementary text detailing the protocol used for enrichment of neural crest cells by HNK-1 labelling and fluorescence activated cell sorting

**Supplementary figures**

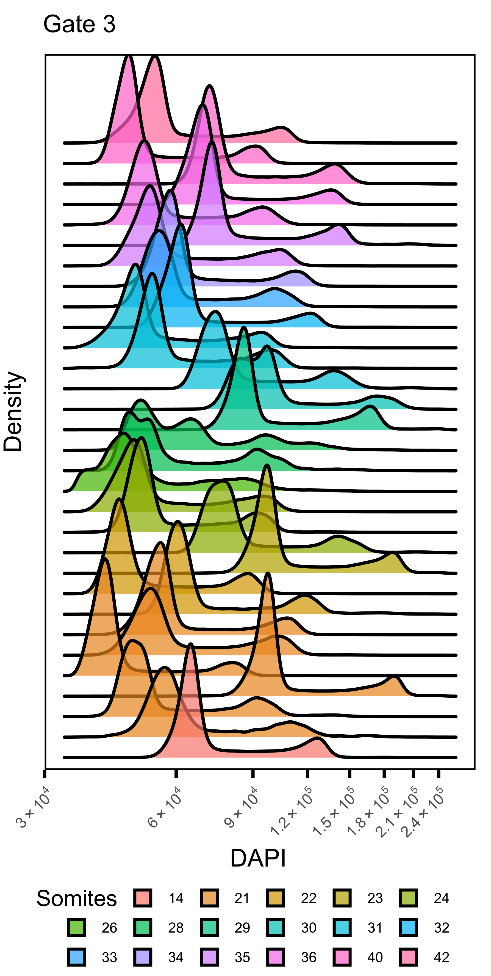

**Supplementary figure 1. Distribution of DAPI-signals across 31 samples subjected to flow cytometry.** In the datasets sorted to gate 3 (see main text), the distribution of DAPI signals revealed typical cell cycles with clearly discernible G1 (2n) and G2 (4n) phases across experiments.

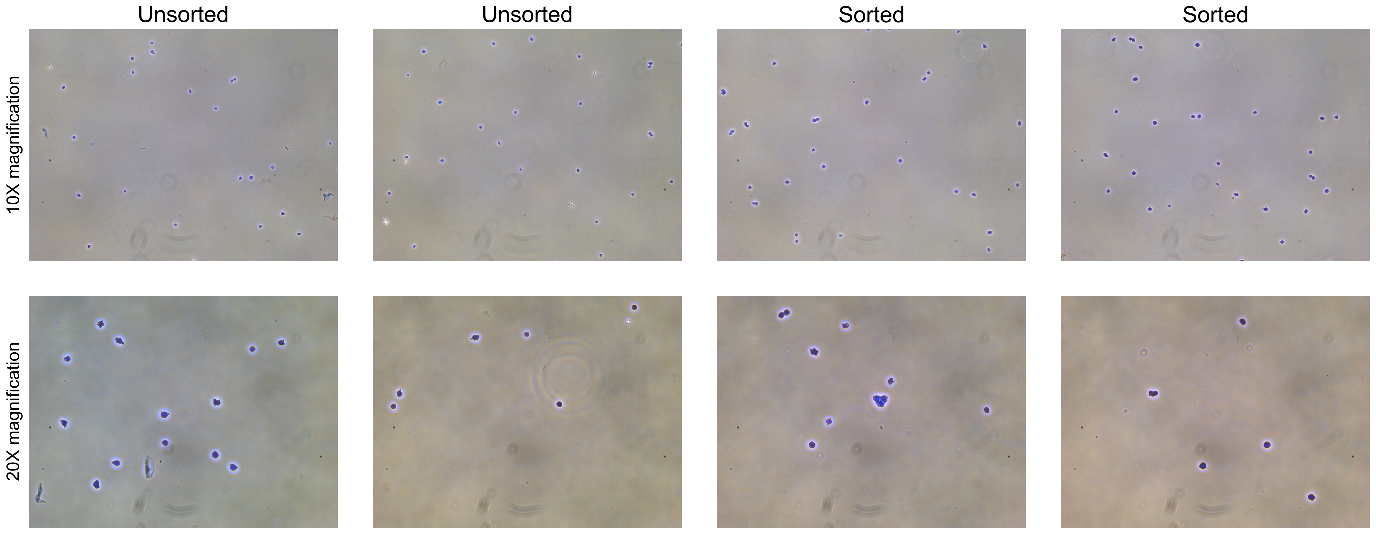

**Supplementary figure 2**. **Light microscopy images of cells that have been dissociated, fixed in methanol and stored at -80°C.** Cells are derived from pooled samples. Two pools have been FAC-sorted and two are unsorted.

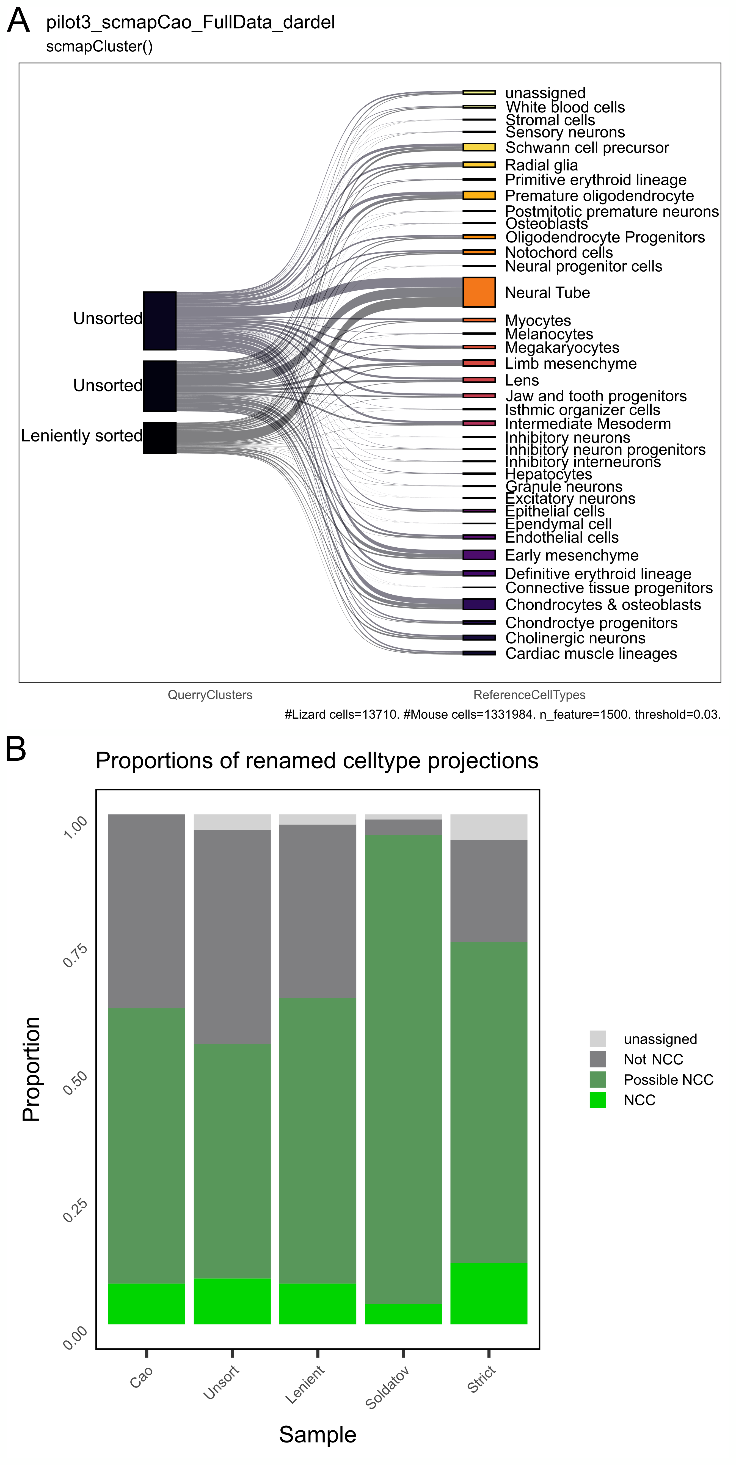

**Supplementary figure 3. A** Projection of the cells from the unsorted and leniently sorted samples onto the whole embryo mouse cell atlas (Cao et al., 2019). **B** The proportion of cells mapping to cell types classified as NCC, not NCC, possible NCC or unassigned. The first bar is the proportion of cell types in the reference dataset (Cao et al., 2019).

**Supplementary tables**

**Supplementary table 1. Cells, reads and gene counts per sample before and after filtering.** Values are provided for the raw data and for the data after filtering (see Material and Methods for details).

|  |  |  | Unsorted | Leniently sorted | Strictly sorted |
| --- | --- | --- | --- | --- | --- |
| Raw data | Cells | n | 15413 | 3382 | 2292 |
|  | Reads/cell | mean | 4153 | 4007 | 44855 |
|  |  | sd | 3505 | 1952 | 20886 |
|  | Genes/cell | mean | 1393 | 1565 | 6217 |
|  |  | sd | 794 | 567 | 1827 |
| Filtered data | Cells | n | 10674 | 3036 | 694 |
|  | Reads/cell | mean | 5156 | 4274 | 45287 |
|  |  | sd | 2804 | 1791 | 13424 |
|  | Genes/cell | mean | 1733 | 1654 | 6737 |
|  |  | sd | 603 | 500 | 1075 |

**Supplementary table 2. Reagent list**

| **Reagent** | **Company** | **Catalog#** | **Cas#** |
| --- | --- | --- | --- |
| PBS | Sigma | P4417 |  |
| Water, nucl. free, Mol. Biol. Grade, Ultrapure | ThermoScientific | J71786.K8 |  |
| Methanol | Merck Millipore | 1.06009.1000 | 67-56-1 |
| Tween-20 | Sigma | P9416 |  |
| Sodium chloride, molecular biology grade (NaCl) | MP | 194848 | 7647-14-5 |
| Bovine serum albumin (BSA) | Sigma | A9418 |  |
| Primary antibody: CD57 monoclonal antibody (TB01 [TBO1]), FITC | eBioscience | 11-0577-42 |  |
| Secondary antibody: Alexa Fluor 488 goat antimouse IgM (u chain) | Invitrogen | A21042 |  |
| DAPI | ThermoScientific | 62248 |  |
| RapiClear 1.49 | SUNJin Lab | RC149001 |  |
| Cover glass 1.5 (~0.17 μm) | VWR | 631-0134 |  |
| iSpacer | SUNJin Lab |  |  |
| RNeasy Micro Kit | Qiagen | 74004 |  |
| SupersScript III reverse transcriptase | Invitrogen | 18080 |  |
| 5x First strand buffer | Invitrogen | Y02321 |  |
| RNase H | New England Biolabs | M0297S |  |
| ROX | Invitrogen | 54881 |  |
| Platinum SYBR Green qPCR SuperMix-UDG | Invitrogen | 11733-046 |  |
| Protein low bind tubes | Fisher | 90410 |  |
| TrypLE Express (1X) Stable Trypsin Replacement Enzyme | Gibco | 12605028 |  |
| Defined Trypsin Inhibitor | Gibco | R-007-100 |  |
| Flowmi^TM^ Cell Strainer |  |  |  |
| LIVE/DEAD^TM^ Fixable Near-IR Dead Cell Stain Kit | Invitrogen | L34975 |  |
| RNasin Plus RNase Inhibitor | Promega | N261A |  |
| MOPS | Sigma | M3183 | 1132-61-2 |
| EGTA | Sigma | E3889 | 67-42-5 |
| Saponin | Sigma | 47036 | 8047-15-2 |
| MgSO_4_ | Sigma | 746452 | 7487-88-9 |
| 1,4-Ditiotreitol (DTT) | VWR | 443852A | 3483-12-3 |

**Supplementary table 3. Primer sequences used in RT-qPCR.**

|  | Forward primer | Reverse primer |
| --- | --- | --- |
| *FoxD3* | AAGAGCAGCTTGGTGAAG | TTGACGAAGCAGTCGTTG |
| *Snai2* | TCAACTTTCTCTGGACTG | AGTAACCATGGTCTAGAG |
| *Sox10* | TGTGAAAGGAGAACAGAG | TGTTCCTTCTTCACCTTC |
| *GAPDH* | TCACAATTTCCAGAGACG | AACAACGTATTGTGCACC |

**Supplementary text**

**Cell dissociation, fixation, and staining – detailed protocol**

Dissect and mince the embryo

1. Place an egg in a sterile petri-dish with nuclease-free phosphate buffered saline (PBS). Open the egg and separate the embryo from the yolk and the surrounding membranes. Transfer the embryo to a new sterile petri-dish with fresh nuclease-free PBS. Take photograph of the embryo to document morphology and developmental stage.
2. Transfer the embryo with a small drop of nuclease-free PBS to a dry sterile petri-dish. Mince the embryo into fine pieces using a sterile scalpel. Work with straight vertical cuts, avoid all grinding and mashing as this will destroy the cells.
3. Use a pasteur pipet to transfer the droplet with the minced embryo to a microcentrifuge tube^[1]^ (1.5 ml). Add more PBS to the remaining parts of the minced embryo and transfer that as well to the tube until all minced pieces of the embryo are transferred. Place the tube on ice.

Dissociate and fix the cells

4. Centrifuge^[2]^ and remove supernatant^[3]^ using a 1 mL micropipette^[4]^ and put the tube back on ice.
5. Resuspend the pellet in 750 µL Trypsin, triturate up and down a few times to make sure the pellet is dissolved in the suspension. Incubate the sample in a water bath for 10 min at 37°C.
6. Add 750 µL Trypsin Inhibitor and immediately triturate up and down 50 times with a 1 mL tip to dissociate the cells. (If there are still visible clumps of cells, centrifuge, remove supernatant, add 1mL nuclease-free PBS, centrifuge and remove supernatant again. Then rerun the Trypsin treatment from step 5 above).
7. Pass the cells through a 70 µm cell strainer^[5]^, centrifuge and resuspend in 500 µL Live/Dead stain (1:1000 in nuclease-free PBS). Incubate on ice for 12 min.
8. Centrifuge, remove supernatant, resuspend in 1000 µL nuclease-free PBS, mix by gentle ttrituration, centrifuge again and remove the supernatant.
9. Resuspend in 30 µL nuclease-free PBS, dissolve the pellet by gentle trituration.
10. Add 1000 µL ice cold methanol. Mix by gently flicking the bottom of the tube with a fingertip until the solution is homogeneous and incubate on ice for 10min. Store the dissociated and fixed cells at -80°C until the day of the flow cytometry experiment.

Label the cells for flow cytometry

11. Centrifuge and remove the supernatant. Resuspend in 300 µL PB and incubate on ice for 5min.
12. Centrifuge and remove the supernatant. Resuspend in 500 µL PB premixed with HNK-1 (1:100) and DAPI (1:1000). Incubate on ice in darkness for 60 min.
13. Centrifuge and remove the supernatant. Resuspend in 300 µL MRDB.
14. Centrifuge and remove the supernatant. Resuspend in 400 µL MRDB and pass through a 70 µm cell strainer.
15. Bring the cells to the flow cytometer and run the experiment.

Buffers:

MRD

0.1M MOPS pH 7.5; 1mM EGTA; 2mM MgSO4; 125mM NaCl; 100U/ml RNAse-inhibitor; 5mM DTT

Do not use DEPC treated water: traces might kill the RNAse inhibitor; filter 0.22µm; make fresh

MRDB

MRD + 2% BSA

PB

MRDB + 0.1% saponin

1. Throughout the protocol, cells should be handled in protein low bind tubes.
2. All centrifugations should be done in a swing centrifuge for 2 min. at 500g and 4°C.
3. Throughout the protocol, the pellet is extremely sensitive. To avoid disturbing the pellet, it is therefore recommended to remove the supernatant directly when the tube is taken from the centrifuge and before placing the tube back on ice. To avoid losing too many cells, it may be preferable to leave some supernatant rather than to remove too much of it.
4. Throughout the protocol, use filter tips.
5. The volue is now >1mL and hence does not fit in a regular 1mL micro pipet tip. Draw the volume in two steps and reuse the same cell strainer for both half of the sample.
